# Experimental reproduction of viral replication and disease in dairy calves and lactating cows inoculated with highly pathogenic avian influenza H5N1 clade 2.3.4.4b

**DOI:** 10.1101/2024.07.12.603337

**Authors:** Amy L. Baker, Bailey Arruda, Mitchell V. Palmer, Paola Boggiatto, Kaitlyn Sarlo Davila, Alexandra Buckley, Giovana Ciacci Zanella, Celeste A. Snyder, Tavis K. Anderson, Carl Hutter, Thao-Quyen Nguyen, Alexey Markin, Kristina Lantz, Erin A. Posey, Mia Kim Torchetti, Suelee Robbe-Austerman, Drew R. Magstadt, Patrick J. Gorden

**Affiliations:** Virus and Prion Research Unit, National Animal Disease Center, Agricultural Research Service, United States Department of Agriculture; Ames, Iowa, 50010, USA; National Veterinary Services Laboratories, Animal and Plant Health Inspection Service, United States Department of Agriculture; Ames, Iowa, 50010, USA; Veterinary Diagnostic and Production Animal Medicine, Iowa State University, Ames, Iowa, 50010, USA

**Keywords:** Highly pathogenic avian influenza, H5N1, clade 2.3.4.4b, genotype B3.13, dairy cattle, mastitis

## Abstract

Highly pathogenic avian influenza (HPAI) H5N1 of the hemagglutinin clade 2.3.4.4b was detected in the United States in late 2021 and continues to circulate in all four North American flyways to date. In addition to impacting poultry, these HPAI viruses caused mortality events in wild bird species and wild mammals. Transmission in multiple host species raises the concern for mammalian adaptation. On March 25, 2024, HPAI H5N1 clade 2.3.4.4b was confirmed in a dairy cow in Texas in response to a multi-state investigation into milk production losses. Over one hundred positive herds were rapidly identified in Texas and eleven other U.S. states. The case description included reduced feed intake and rumen motility in lactating cows, decreased milk production, and thick yellow milk. The diagnostic investigation revealed detections of viral RNA in milk and mammary tissue with alveolar epithelial degeneration and necrosis, and positive immunoreactivity of glandular epithelium by immunohistochemistry. A single transmission event, likely from avian species to dairy cattle, followed by limited local transmission preceded the onward lateral transmission of H5N1 clade 2.3.4.4b genotype B3.13. We sought to experimentally reproduce infection with genotype B3.13 in Holstein yearling heifers and lactating cows. The heifers were inoculated by an aerosol respiratory route and the cows by an intramammary route. Clinical disease was mild in the heifers, but infection was confirmed by virus detection, lesions, and seroconversion. Clinical disease in lactating cows included decreased rumen motility, changes to milk appearance, and production losses consistent with field reports of viral mastitis. Infection was confirmed by high levels of viral RNA detected in milk, virus isolation, lesions in mammary tissue, and seroconversion. This study provides the foundation to investigate additional routes of infection, transmission, and intervention strategies.

## Introduction

Highly pathogenic avian influenza (HPAI) H5N1 viruses of the goose/Guangdong lineage in the hemagglutinin clade 2.3.4.4b were detected in the United States (U.S.) in 2021 following widespread dispersion across Asia and Europe. H5N1 in this clade were detected across the U.S. in mortality events in wild bird species and wild mammals. Additionally, outbreaks occurred in numerous commercial and backyard poultry premises, leading to culling to control the spread. On March 25, 2024, the US Department of Agriculture (USDA) National Veterinary Services Laboratories (NVSL) confirmed a case of HPAI H5N1 clade 2.3.4.4b genotype B3.13 in a dairy cow in Texas in response to a multi-state investigation into milk production losses. Additional outbreaks were rapidly identified in Texas as well as eight other U.S. states. As of July 11, 2024, 146 cases were confirmed in 12 states ^1^. The typical case description on the affected dairy farms included reduced feed intake and rumination in lactating cows, rapid drop in milk production, and affected cows with thick yellow milk with flecks and/or clots ^2^. The diagnostic investigation revealed low Ct detections of viral RNA in RT-qPCR assays, mammary tissue with alveolar epithelial degeneration and necrosis with intraluminal sloughing of cellular debris, and positive immunoreactivity of glandular epithelium with nuclear and cytoplasmic labeling by immunohistochemistry (IHC). Deceased domestic cats that presented with neurologic and respiratory signs from the affected dairy herds were also diagnosed and confirmed to be infected with the same H5N1 clade 2.4.4.4b B3.13 genotype, as well as several peridomestic wild birds found around or near affected premises.

A genomic and epidemiologic investigation demonstrated that a single transmission event from avian species preceded the multi-state cattle outbreak ^3^. The movement of preclinical or subclinical lactating dairy cows was the major contributor to spread among the U.S. dairy premises. Sequence analysis thus far revealed highly similar genomes among cattle detections with little evolution or known mammalian adaptation markers ^3^. Several human infections were reported with H5N1 clade 2.3.4.4b, and although some were severe or fatal, the four human detections associated with the dairy outbreak in the U.S. were mild ^4–6^. However, transmission in multiple host species, particularly mammals, raises the concern for mammalian adaptation that may lead to increased potential for human infection and/or transmission.

Although a few sporadic detections of human seasonal influenza A virus (IAV) or antibodies were previously reported in dairy cows associated with milk production loss ^7,8^, the Texas cases in 2024 were the first reports of HPAI of any subtype causing viral mastitis in lactating dairy cows. At the time we initiated these experiments, the route of infection and transmission between cows was unknown. Transmission between farms was linked to movement of live lactating cows, yet within farm spread to resident cows was observed within days or weeks following movement without a clear pattern of transmission consistent on all farms. Here, we sought to experimentally reproduce infection of dairy cattle with genotype B3.13 in Holstein yearling heifers via an aerosol respiratory route and in lactating cows via an intramammary route.

## Materials and Methods

### Strain characterization

We analyzed the HPAI H5N1 genotype B3.13 virus used this study (A/dairy cattle/Texas/24-008749-002/2024: TX/24: NCBI PP755581-PP755588) with other B3.13 strains generated within Nguyen et al.^3^ and newly sequenced data collected between April 16 and May 8, 2024. Influenza A virus RNA from samples was amplified, then we generated cDNA libraries by using the Illumina DNA Sample Preparation Kit, (M) Tagmentation (Illumina, https://www.illumina.com) and either the 300 or 500-cycle iSeq or NextSeq Reagent Kit v2 (Illumina) according to manufacturer instructions. We performed reference-guided assembly of genome sequences using IRMA v1.1.5 ^9^. We aligned each gene segment using mafft v7.490 ^10^ and inferred maximum likelihood phylogenetic trees for each gene segment as well as concatenated whole genomes using IQ-Tree v2.2.2 ^11^. These phylogenetic gene trees were used to determine how representative the TX/24 strain was relative to viruses collected between March and May 2024. This approach implemented the evaluate algorithm within PARNAS ^12^ that objectively identified the most representative strain within the B3.13 group and how near the TX/24 strain was relative to that representative strain.

### Cow intramammary inoculation

All animal studies were carried out in a BSL3-Ag facility in compliance with the Institutional Animal Care and Use Committee of the USDA-ARS National Animal Disease Center (NADC) and Federal Select Agent Program regulations. Two non-pregnant, lactating Holstein cows of approximately 3 years of age and 280 days in milk during their first lactation were moved from the NADC dairy into BSL3-Ag containment and acclimated for approximately 1 week before inoculation. Cows were inoculated with 1 ml of 1 x 10^5^ tissue culture infectious dose 50 (TCID_50_)/ml A/dairy cattle/Texas/24-008749-002/2024 by an intramammary route in each of two quarters, front right and rear left, using a teat canula (Supplemental Figure 1A). After instilling, the inoculum was moved upward into the teat sinus with gentle massage.

### Calf aerosol inoculation

Five Holstein heifer calves of approximately one-year of age were inoculated with either 2 ml of 1 x 10^6^ TCID_50_/ml A/dairy cattle/Texas/24-008749-002/2024 (n=4) or sham-inoculated with phosphate buffered saline (PBS) (n=1) by an aerosol respiratory route. For aerosol inoculation, calves were restrained in a cattle stanchion and the inoculum delivered by nebulization into a mask covering the nostrils and mouth (Supplemental Figure 1B). Two ml of inoculum were placed into the medicine cup of the portable equine nebulizer (Flexineb E3, Nortev Ltd, Oranmore, Galway, Ireland). Nebulization continued until all the inoculum was delivered (approximately 3 min). Following the inoculum, 2 ml sterile PBS were nebulized through the apparatus. The portable equine nebulizer uses a vibrating mesh nebulizer, which generates aerosol droplets in the respirable range. Droplets generated by vibrating mesh nebulizers were shown to be 3-5 um ^13^, with the manufacturer of the nebulizer reporting that 70% of the total nebulized volume resulted in droplets of <5 um. Aerosolized droplets that are less than 5 um bypass the upper respiratory tract and are deposited deep in the lower respiratory tract ^14^.

### Clinical evaluation and sample collection

In the lactating cows, rumination and other behaviors were monitored continuously using an ear-tag accelerometer sensor (CowManager SensOor, Agis Automatisering BV, Harmelen, the Netherlands). For cows and heifers, clinical signs were visually monitored and recorded daily (Supplemental Table 1), including body temperature with a thermal microchip, respiratory effort and rate, nasal and ocular discharge, and fecal consistency. The lactating cows were milked once daily in the morning using a portable milker (Hamby, Maysville, MO). Prior to milking, the teats were cleaned with water, disposable paper towels, and isopropyl alcohol wipes, and milk manually stripped from each teat into separate 50 ml tubes. The milk samples were evaluated by quarter for mastitis by California Mastitis Test (CMT, ImmuCell, Portland, ME) and evaluation for consistency and color using a dental color scorecard on a scale of 0-12. The remaining milk stripped by quarter was used for virus and/or antibody detection. After milking, a sample from each cow’s milking machine bucket was also collected. A separate milking claw was used for each cow, and the claw, tubing, and bucket were washed, degreased, and disinfected (Virkon S, LanXess, Pittsburg, PA) after each use.

The remaining clinical samples were collected daily from cows and calves for 7 days post-inoculation (DPI) and approximately every 2 days thereafter until study termination. To sample the deep pharyngeal region, a Frick cattle mouth speculum was used. Using manual restraint, the speculum was placed over the tongue to the pharynx and held there. Through the speculum, FLOQ nylon-tipped (Copan, Murrieta, CA) or long cotton-tipped (SCA Health, Dallas, TX) swabs were used to collect samples of the deep oropharynx. Ocular, nasal, and rectal samples were collected with FLOQ nylon swabs. Whole blood was collected via jugular venipuncture and placed into molecular transport media (PrimeStore MTM, Longhorn, Bethesda, MD) and into serum separator tubes. Saliva was collected with an absorbent pad (Super-SAL, Oasis Diagnostics, Vancouver, WA).

### Pathologic evaluation

All cattle were humanely euthanized via intravenous administration of pentobarbital sodium (Fatal Plus, Vortech Pharmaceuticals, Dearborn, MI). Heifers 2311 and 2316 and a negative control heifer were necropsied at 7 DPI while the remaining two aerosol inoculated heifers were necropsied at 20 DPI. The lactating dairy cows were necropsied at 24 DPI. At the time of necropsy, the thoracic cavity, abdominal cavity and cranium (longitudinal section) underwent macroscopic evaluation. Paired fresh and formalin-fixed tissues for RT-qPCR and microscopic evaluation, respectively, were collected (Supplemental Table 2). Formalin-fixed tissues were processed routinely for microscopic evaluation. Additional fresh samples included aqueous and vitreous humor, urine, feces and rumen contents.

Immunohistochemistry (IHC) targeting the nucleoprotein (NP) antigen of IAV was performed as previously described with the inclusion of a positive and negative tissue control ^15^. A representative section was selected for IHC from nasal turbinate, trachea, lung, tracheobronchial lymph node, supramammary lymph node (lactating cows), and individual mammary quarters (lactating cows) from each animal. Additional tissue sections with RT-qPCR detection and/or histologic lesions were also evaluated by IHC for the detection of NP antigen to confirm a causal link between IAV and the lesion. A Masson’s trichrome stain was performed per the manufacturer’s instructions on representative mammary gland sections from each of the four quarters to highlight fibrous connective tissue (Newcomer Supply).

### Virus detection in clinical samples

IAV RNA extraction and reverse transcription quantitative real-time PCR (RT-qPCR) were performed at the USDA NVSL according to the standard operating procedures ^16^. Clinical ante-mortem and necropsy samples were tested using an IAV matrix gene RT-qPCR. A subset of positive samples with Ct ≤35 was processed for whole genome sequencing (WGS). Influenza A virus RNA from samples was amplified ^17^; and after amplification was completed, we generated cDNA libraries for MiSeq as indicated under virus characterization. A subset of RT-qPCR positive samples was also inoculated onto 10-day old embryonating chicken eggs for virus isolation to determine the presence of viable virus.

### Antibody Detection

Seroconversion was determined using a blocking ELISA to detect antibodies to the nucleoprotein (NP) (Influenza A Ab, IDEXX, Westbrook, Maine) according to the manufacturer instructions for swine at the time of the study with a 1:10 starting dilution of serum or rennet-treated milk and a cut-off sample to negative (S/N) optical density (O.D.) ratio of 0.6. Prior to ELISA, milk samples were treated with rennet (Mucor miehei, Sigma-Aldrich, St. Louis, MO) as previously described^18^. H5 specific hemagglutinin inhibition (HI) and virus neutralizing (VN) antibody assays on the London line of Madin-Darby Canine Kidney (MDCK) cells were conducted as previously described^19–21^. Prior to HI, sera were heat inactivated at 56°C for 30 minutes, treated with receptor destroying enzyme (Hardy Diagnostics, Santa Maria, CA), and adsorbed with 0.5% and subsequently 100% rooster red blood cells for 20 minutes each to remove nonspecific hemagglutinin inhibitors and natural serum agglutinins.

### Processing of raw sequence data and single nucleotide variant calling

To analyze the Illumina short read data for 82 samples, we used the “Flumina” pipeline (https://github.com/flu-crew/Flumina) for processing and analyzing influenza data ^3^. The pipeline uses Python v3.10, R v4.4 (R Development Core Team 2024), and SnakeMake ^22^ to organize programs and script execution. The reads are preprocessed using FASTP ^23^, removing adapter contamination, low complexity sequences, and other artifacts. Consensus contigs for phylogenetics were assembled using IRMA v1.1.4 ^9^. The pipeline maps cleaned reads to the reference cattle strain (A/dairy cattle/Texas/24-008749-002/2024) using BWA (bwa index –a bwtsw;) ^24^. High frequency single nucleotide variants (SNV) were called with GATK v.4.4 ^25^ and low frequency SNVs were called with LoFreq ^26^. To assess potential SNV phenotypic changes, a database was generated using the Sequence Feature Variant Types tool from the Influenza Research Database ^27^ for all eight genes. To estimate genome-wide natural selection, we used the program SNPGenie on the VCF files ^28^.

### Statistical Analysis

Pearson correlation coefficients between Ct values for milk samples from the bucket as well as the inoculated teats (front right (FR) and rear left (RL)) and rumination time, milk production and consistency and color scores of milk from the inoculated teats were calculating using the package Hmisc v. 5.1-3c in RStudio 2022.02.1+461 “Prairie Trillium”. Correlations were considered significant at *P* < 0.05 and visualized using the package corrplot v. 0.92.

## Results

### Phylogenetic analysis of challenge strain

The phylogenetic tree topology (Figure 1) with B3.13 genotype strains collected from dairy cattle between March to May 2024 was congruent with earlier analyses ^3^. The HA gene tree indicated a single monophyletic clade; following the introduction of the B3.13 virus genotype into dairy cattle in late 2023 ^3^, it persisted and rapidly spread through cattle populations. There was no evidence of reassortment, and all viruses detected in dairy cattle were classified as B3.13 genotype maintaining the new PB2 and NP gene that had been acquired in migratory birds prior to the spillover ^3^. The HA gene tree and the phylogenomic tree had little evidence for directional selection with long external branches relative to internal branches, and many viruses that were identical. These data suggested a founder effect where a single genotype was introduced into a new population of susceptible hosts ^29^. The TX/24 challenge strain was 99.94% identical to a B3.13 whole genome consensus strain (9 nucleotide substitutions across the genome); and at the amino acid level, the TX/24 HA gene was identical to a consensus B3.13 HA gene sequence with only two synonymous substitutions detected.

**Figure 1.**
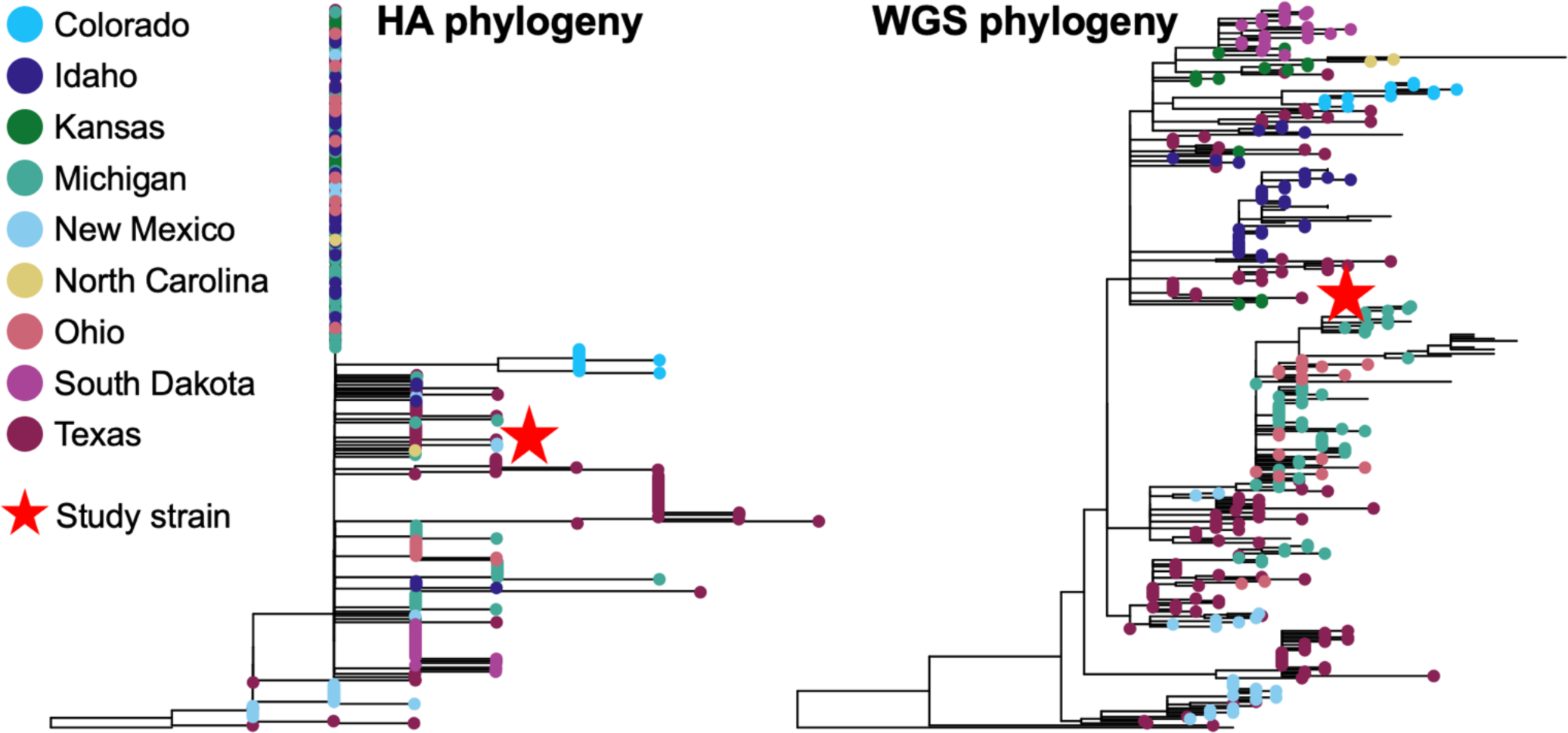
Evolutionary history of H5N1 clade 2.3.4.4b hemagglutinin genes from dairy cattle in the United States collected between March and May 2024. The HA gene and whole genome phylogeny demonstrated a single introduction of the virus from wild birds to dairy cattle with subsequent dissemination across nine US states. The A/dairy cattle/Texas/24-008749-002/2024 study strain, indicated by a red star, was 99.94% identical to a B3.13 whole genome consensus strain (9 nucleotide substitutions across the genome); and at the amino acid level, the TX/24 HA gene was identical to a consensus B3.13 HA gene sequence with only 2 synonymous substitutions detected.

### Clinical observations

Following transition to containment, rumen motility, feed consumption, and milk production stabilized in the lactating cows by the time of inoculation. Following inoculation, both cows showed signs of mastitis with positive CMT and milk color and consistency changes beginning at 2 DPI and lasting until approximately 14 DPI, only in the inoculated quarters (Figure 2A). The peak average color change score in the milk from inoculated quarters from both cows was 6.5 on a yellow/brown scale (0-12) on 5 and 6 DPI. The milk was yellow, thicker, and contained flakes and clots of debris from affected quarters during the same time period. Rumen motility starkly declined on 1 DPI and began recovering after 7 DPI (Figure 2B). Milk production steadily declined from 1-4 DPI, remained low until 10-12 DPI, and was 71-77% of pre-inoculation production at 23 DPI (Figure 2C). The cows showed signs of lethargy, reduced feed intake, self-resolving watery diarrhea or dry feces, and intermittent clear nasal discharge throughout the 24-day observation period. Neurologic signs were not observed. The heifer calves displayed no overt signs of illness, with only transient increases in nasal secretions observed. Other clinical parameters indicative of illness were not consistently observed in the lactating cows or heifer calves.

**Figure 2.**
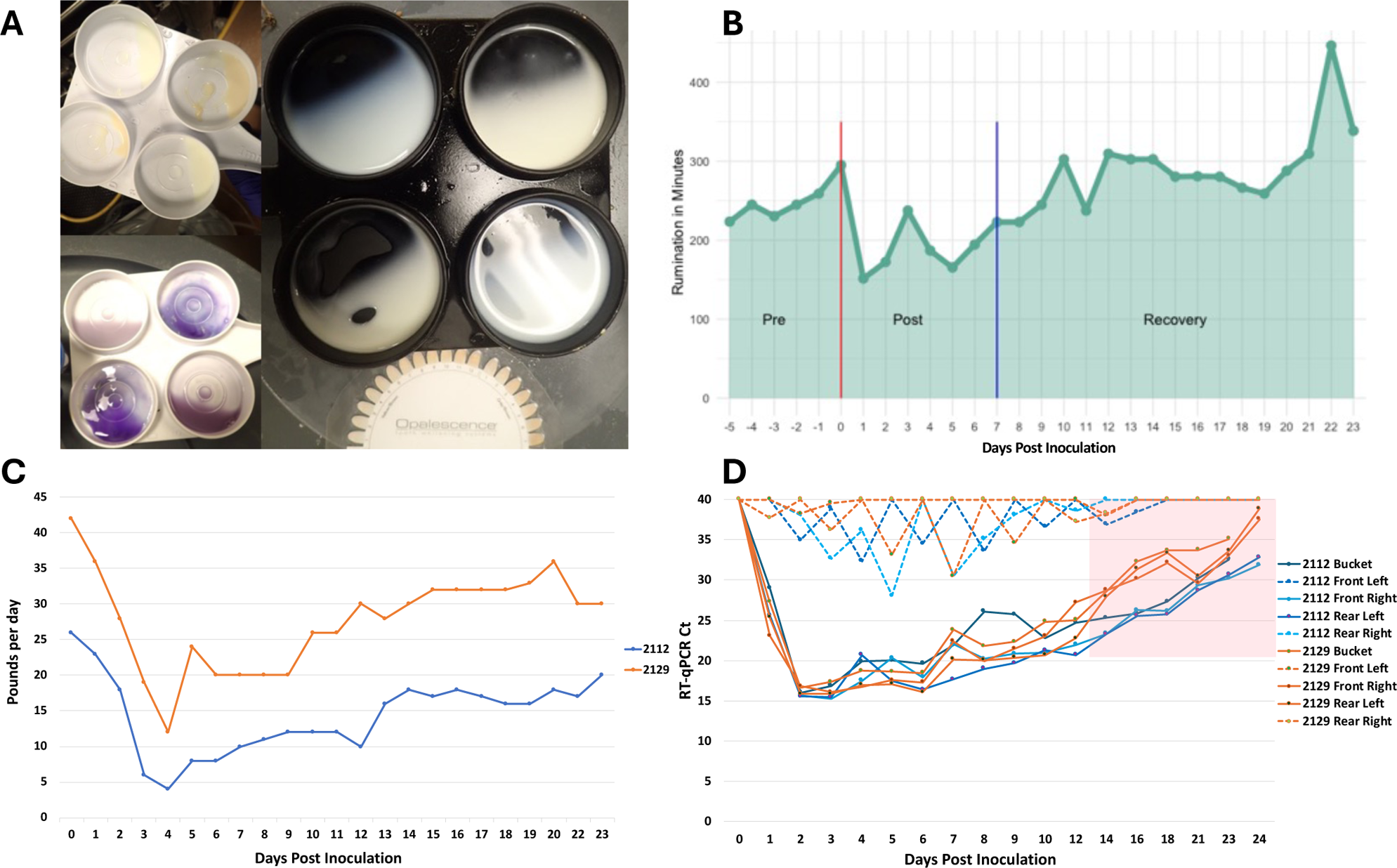
Clinical signs and viral detection in lactating dairy cows. A) Representative milk sampling demonstrating thickening, flakes or clots, and change in color in the upper left and right photographs, and gel formation in positive California Mastitis Tests in the lower left photograph. In all photos, changes were observed in the inoculated front right and rear left quarters in the upper right and lower left cups of the milk collection paddle in each photo. B) Average daily rumination time in minutes for both lactating cows measured using an ear-tag accelerometer sensor. Rumination time decreased at one day post inoculation (DPI) and recovered to pre-challenged levels at 7 DPI. C) Each individual milking machine bucket was weighed once daily to monitor production, Cow 2112 (blue) and Cow 2129 (orange). Milk production steadily declined from 1-4 DPI, remained low until 10-12 DPI, and was 71-77% of pre-inoculation production at 23 DPI. D) Milk was stripped from each teat prior to milking and a sample from the bucket after milking, Cow 2112 (blue) and Cow 2129 (orange). An RT-qPCR assay detected viral RNA beginning on 1 DPI until study termination at 24 DPI in the inoculated quarters (solid lines), with cycle threshold (Ct) on the y-axis and DPI on the x-axis. The positive detections in non-inoculated quarters (dashed lines) were likely cross-contamination due to the non-sterile collection of milk from each teat. The pink shading indicates time points when live virus isolation was attempted from the bucket samples and were negative by egg inoculation. One inoculated quarter from Cow 2112 was VI positive on 12 DPI, but all tested samples beyond 12 DPI were VI negative.

### Virus detection in clinical and necropsy samples

Milk stripped from inoculated mammary quarters and the post-milking bucket sample were continuously RT-qPCR positive from 1 DPI until the end of the study (Figure 2D). Although stripped milk from non-inoculated quarters had Ct below 35, the detection was not consistent over consecutive days and never progressed to low Ct values indicative of live virus replication in those mammary glands. Milk bucket samples subjected to virus isolation were positive on 5, 7, and 10 DPI and negative on 12, 14, 16, and 18 DPI. One inoculated quarter from Cow 2112 was VI positive on 12 DPI, but all tested mammary quarter samples beyond 12 DPI were VI negative.

There were strong positive correlation coefficients between Ct values from the milk bucket and infected quarters with rumination and milk. As Ct decreased (higher viral load) so did rumination and milk production. There were also strong negative correlation coefficients between milk Ct values and color and consistency scoring, as Ct decreased (higher viral load), color and consistency scores increased (Figure 3). Only one nasal swab (Ct 33.1) from Cow 2112 and one ocular swab (Ct 34.7) from Cow 2129 on 3 DPI had RT-qPCR Ct ≤35; all remaining clinical samples had RT-qPCR Ct greater than 35 cycles or undetected. No fecal swab or blood samples were RT-qPCR positive at any time point. From the broad range of samples collected at necropsy, mammary tissue from inoculated quarters and supramammary lymph nodes had RT-qPCR Ct ≤35, an inguinal lymph node and one inoculated mammary tissue had Ct between 35-38. All other samples were not within RT-qPCR detection limits. (Supplemental Table 2).

**Figure 3.**
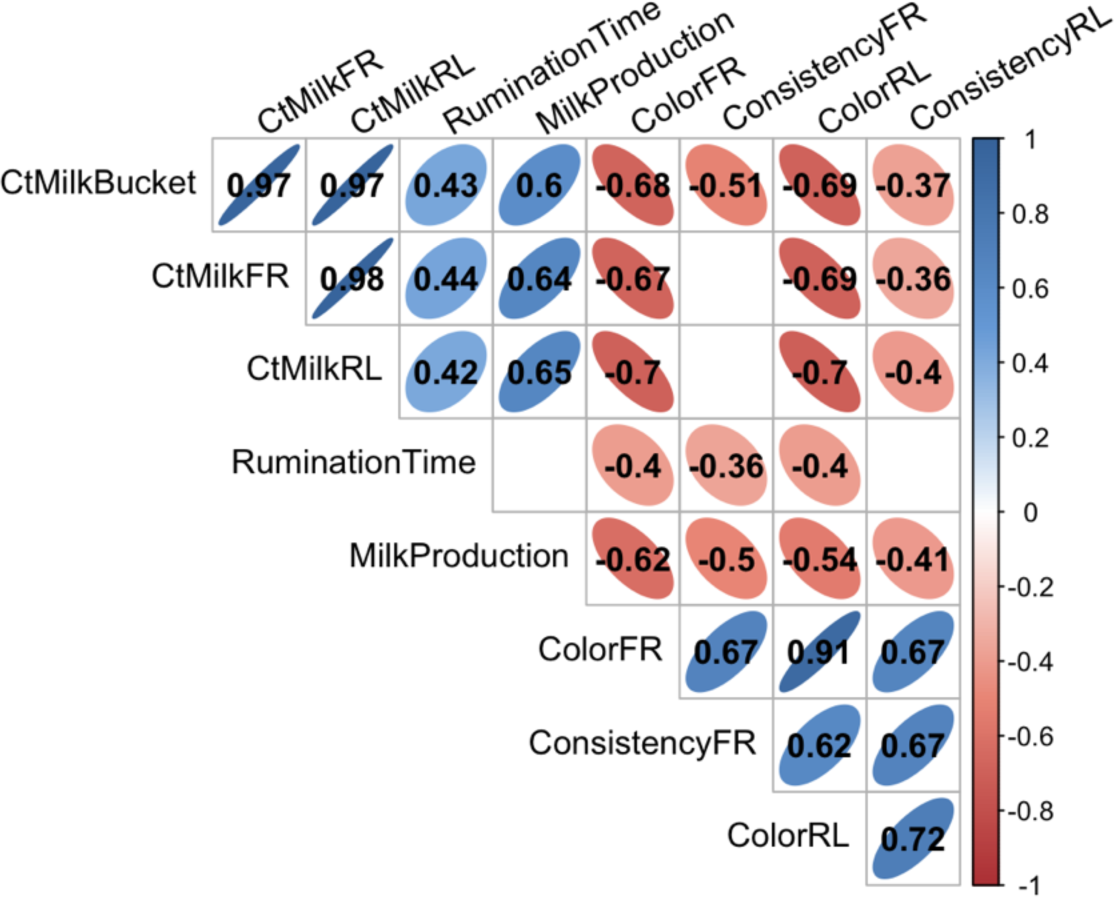
Significant (P<0.05) Pearson correlation coefficients between Ct values for milk samples from the bucket as well as the inoculated teats (front right (FR) and rear left (RL)) and rumination time, milk production and consistency and color scores of milk from the inoculated teats. Empty squares in the matrix indicated the correlation was not significant (P<0.05)

Nasal, oropharyngeal, saliva, and ocular samples collected from the aerosol inoculated heifers sporadically had RT-qPCR Ct ≤35 (Supplemental Table 3). Heifer 2316 had nasal swabs with Ct ≤35 from 1-5 and 7 DPI. Viral RNA was also detected across multiple days in oropharyngeal swabs, ocular swab, or saliva from Heifer 2316. Viable virus isolation was attempted on clinical samples with Ct ≤35 and were VI positive for nasal swabs from Heifer 2316 on 2-5 and 7 DPI. This heifer was euthanized for necropsy on 7 DPI, so there were no further ante-mortem samples collected. Each of the other three heifers tested RT-qPCR positive on at least one time point between one and seven DPI, but sporadically and not on consecutive days. Heifer 2311 had one VI positive FLOQ oropharyngeal swab on 3 DPI. No fecal swab or blood samples were RT-qPCR positive at any time point. The sham-inoculated negative control Heifer 2309 had no RT-qPCR detections as expected. From the broad range of samples collected at the 7 DPI necropsy, lung tissue from both heifers (accessory lobes and cranial part of the left cranial lobe) had RT-qPCR Ct ≤35, and a retropharyngeal lymph node, lung lobe (caudal part of the right cranial lobe and middle lobe), and a turbinate sample had Ct between 35-38. All other samples at DPI 7 as well as all samples from the 20 DPI necropsy were not within RT-qPCR detection limits (Supplemental Table 2).

### Single nucleotide variant analysis of clinical samples

Across 82 samples collected from the experimentally inoculated cattle, we identified within-host SNVs that were present at low frequencies (present in greater than 0.5 percent) across the genome. We matched a custom database of variants of interest that could potentially provide a selective advantage and alter virus phenotype. Overall, there were 3,960 SNVs present at low frequencies, where 2,676 altered the amino acid with nonsynonymous changes (Supplemental Table 4). Of the amino acid changes detected, only 12 had been previously associated with known functional changes. We detected low frequency variants associated with changes in pathogenicity/virulence in MP (T139A), NS (T91N/A, D92E/N, T94A, L95P, S99P, D101N/G, D125N), and PB1 (R622Q). We also detected low frequency variants associated with mammal adaptation in NS (F103S) and PB2 (L631P).

### Antibody responses

The dairy cows inoculated by the intramammary route were negative for NP antibody in serum prior to inoculation, became positive on 7 DPI using a sample/negative (S/N) ratio cut-off of 0.6, and remained positive on all sample timepoints (Table 1A). Fresh stripped milk from both cows were also positive by NP ELISA by 9 DPI. Both cows were seropositive by HI assay on 24 DPI and by VN on 14 and 24 DPI (Table 1B). Milk samples collected from each quarter on 24 DPI were tested by HI with reciprocal titers of 10-40, with both cows having quarters above and below the HI positive cut-off of ≥40. However, VN in inoculated quarters were positive on 9 DPI for Cow 2129 and on 12 DPI for Cow 2112. The uninoculated quarters from both cows remained below the positive cut-off titer of 1:40. Two of the heifer calves were positive for NP antibodies on 7 DPI while the remaining two seroconverted between 9 and 13 DPI (Table 2). One calf was positive by HI assay and the calf with a later NP antibody response was suspect at 20 DPI. The sham inoculated negative control remained seronegative as expected.

**Table 1A.** Serum and milk ELISA and hemagglutination inhibition antibody response in lactating dairy cows.*

| ID | Sample | DPI | 0 | 4 | 5 | 6 | 7 | 8 | 9 | 10 | 12 | 14 | 24 | HI |
| --- | --- | --- | --- | --- | --- | --- | --- | --- | --- | --- | --- | --- | --- | --- |
| 2112 | Serum |  | 0.907 | 1.252 | 1.265 | 0.742 | 0.394 | 0.296 | 0.299 | 0.331 | 0.338 | 0.323 | 0.194 | 80 |
| 2129 | Serum |  | 0.796 | 1.406 | 1.533 | 0.975 | 0.490 | 0.353 | 0.337 | 0.320 | 0.335 | 0.254 | 0.261 | 160 |
| 2112 | Milk |  |  | 1.317 | 1.164 | 1.000 | 0.871 | 0.585 | 0.481 | 0.352 | 0.365 | 0.270 | 0.297 | # |
| 2129 | Milk |  |  | 2.036 | 1.154 | 0.939 | 0.805 | 0.664 | 0.473 | 0.347 | 0.265 | 0.255 | 0.231 | # |

\*NP ELISA sample to negative (S/N) ratio of optical densities reported for days post inoculation (DPI) indicated in header row. Hemagglutination inhibition (HI) assay reported as reciprocal titer on the final timepoint of 24 DPI. Positive samples indicated by gray shading and italics. #ELISA S/N reported for pooled milk bucket samples. HI titers in milk at 24 DPI varied by quarter between 10-40.

**Table 1B.**
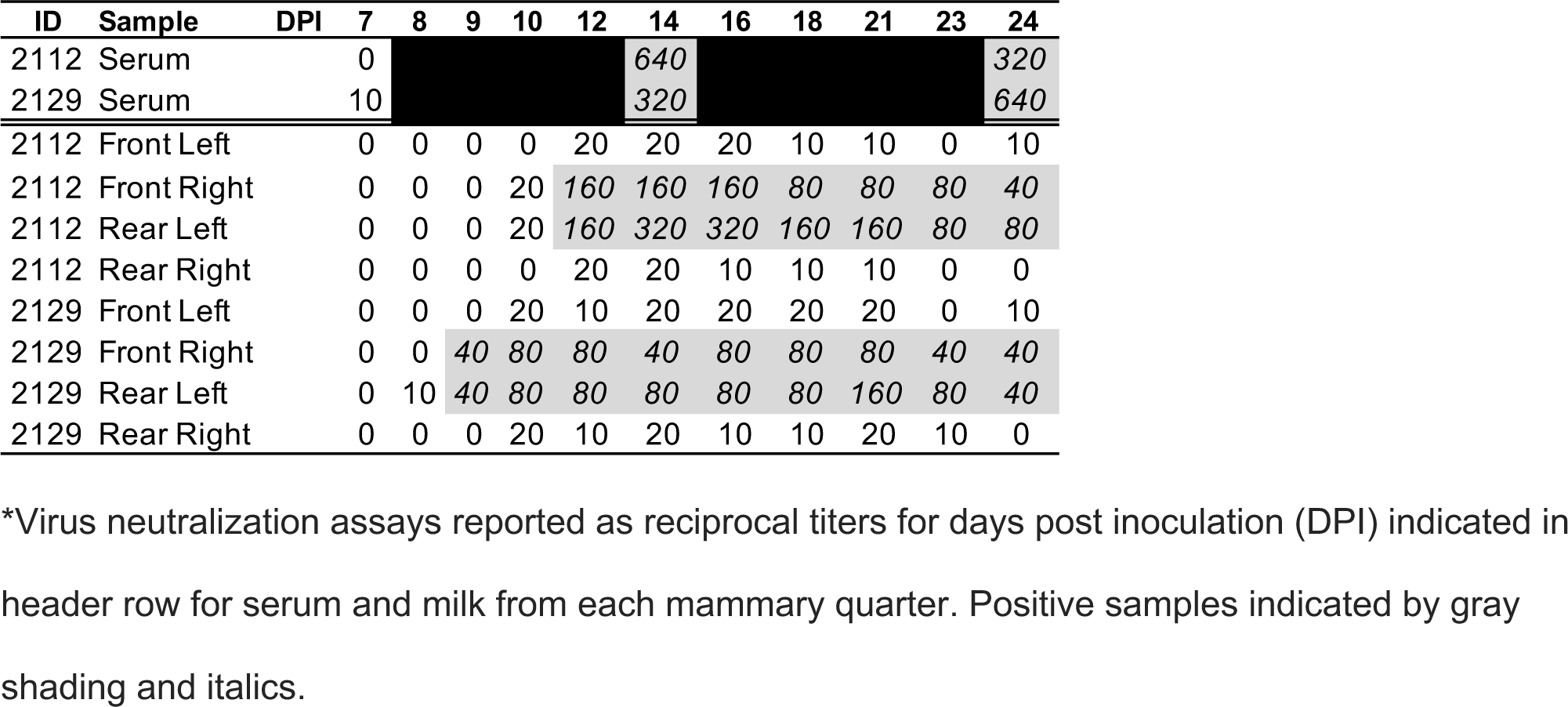
Serum and milk virus neutralizing antibody responses in lactating dairy cows.

**Table 2.** Serum antibody response in yearling heifers.

| ID | DPI | 0 | 3 | 7 | 9 | 13 | 20 | HI | VN |
| --- | --- | --- | --- | --- | --- | --- | --- | --- | --- |
| 2311 |  | 0.923 | 1.252 | 0.455 |  |  |  |  |  |
| 2312 |  | 0.783 | 0.944 | 0.828 | 0.974 | 0.454 | 0.520 | 20 | 20 |
| 2313 |  | 0.895 | 1.186 | 0.900 | 0.401 | 0.387 | 0.370 | 40 | 40 |
| 2316 |  | 0.977 | 1.162 | 0.499 |  |  |  |  |  |

\*NP ELISA sample to negative (S/N) ratio of optical densities reported for days post inoculation (DPI) indicated in header row. Hemagglutination inhibition (HI) and virus neutralization (VN) assays reported as reciprocal titer on the final timepoint of 20 DPI. Positive samples indicated by gray shading and italics. Two animals were euthanized on 7 DPI.

### Pathologic evaluation

Minimal multifocal pulmonary consolidation was present in Heifer 2316 (Figure 4). In the lactating cows, lesions considered incidental were observed and included a unilateral locally extensive pleural adhesion and transparent, straw-colored abdominal fluid in cow 2112 and interlobular pulmonary edema in cow 2129. Macroscopic evaluation of the remaining tissues was unremarkable.

**Figure 4.**
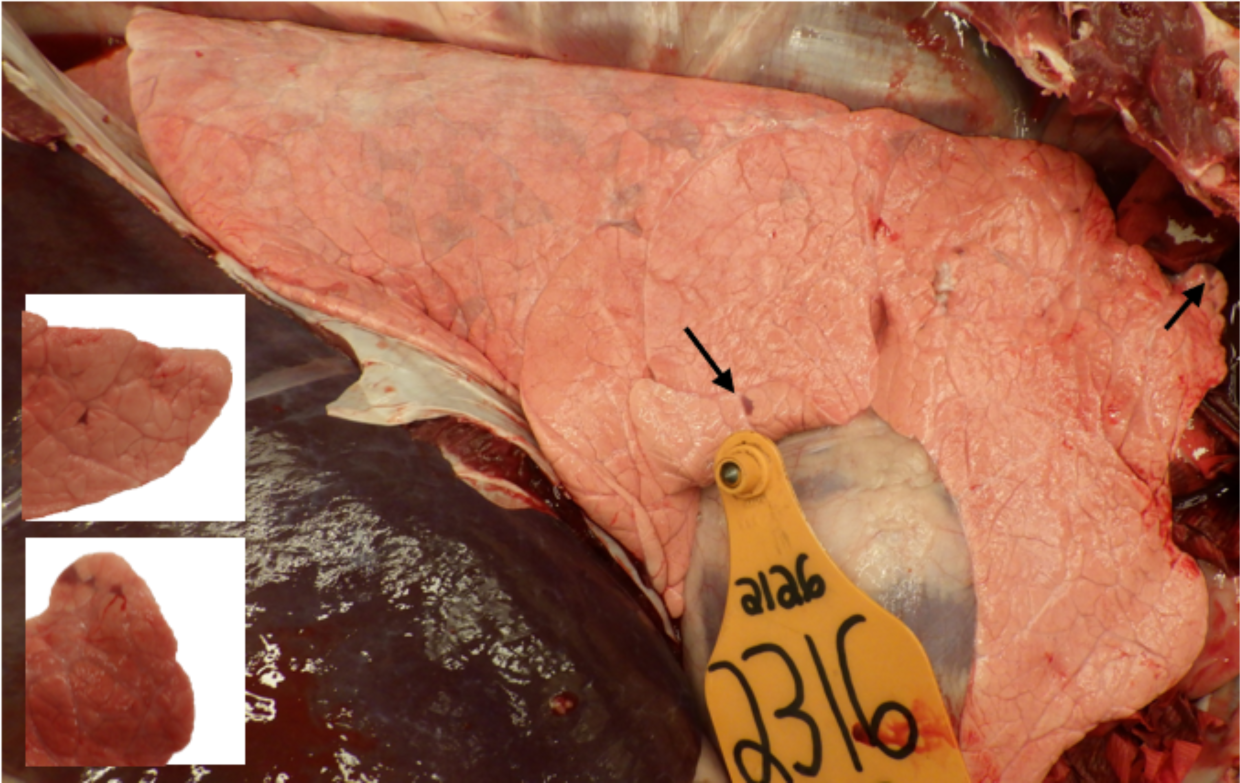
Minimal multifocal pulmonary consolidation (arrows and insets) consistent with influenza A virus infection in Heifer 2316.

Similar histologic lesions were present in the left rear mammary gland (inoculated) of both dairy cattle. Fibrous connective tissue replaced 15% to 50% of secretory alveoli and ducts, both intralobular and interlobular, of multifocal lobules (Figure 5A-D). The remaining secretory alveoli in affected lobules were commonly shrunken and lined by small, attenuated, or swollen, vacuolated epithelial cells (atrophy and degeneration). Both interlobular and intralobular fibrous connective tissue contained multifocal aggregates of segmental to circumferential perialveolar or periductular mononuclear inflammatory cells predominated by lymphocytes and plasma cells. The remaining secretory alveoli and ducts contained secretory product admixed with occasional foamy cells (macrophages or foam cells) or rarely deeply basophilic, concentrically lamellated foci (corpora amylacea). Nonaffected to minimally affected glandular tissue commonly contained moderately to well-developed secretory alveoli. Histologic evaluation of the non-inoculated mammary quarters (left front and right rear) was characterized by well-developed secretory epithelium that contained fluid and no to minimal intra- or interlobular fibrosis and no to rare aggregates of mononuclear leukocytes predominated by lymphocytes and plasma cells (Figure 5E and F). A summary of microscopic findings including IHC is presented in the data file.

**Figure 5.**
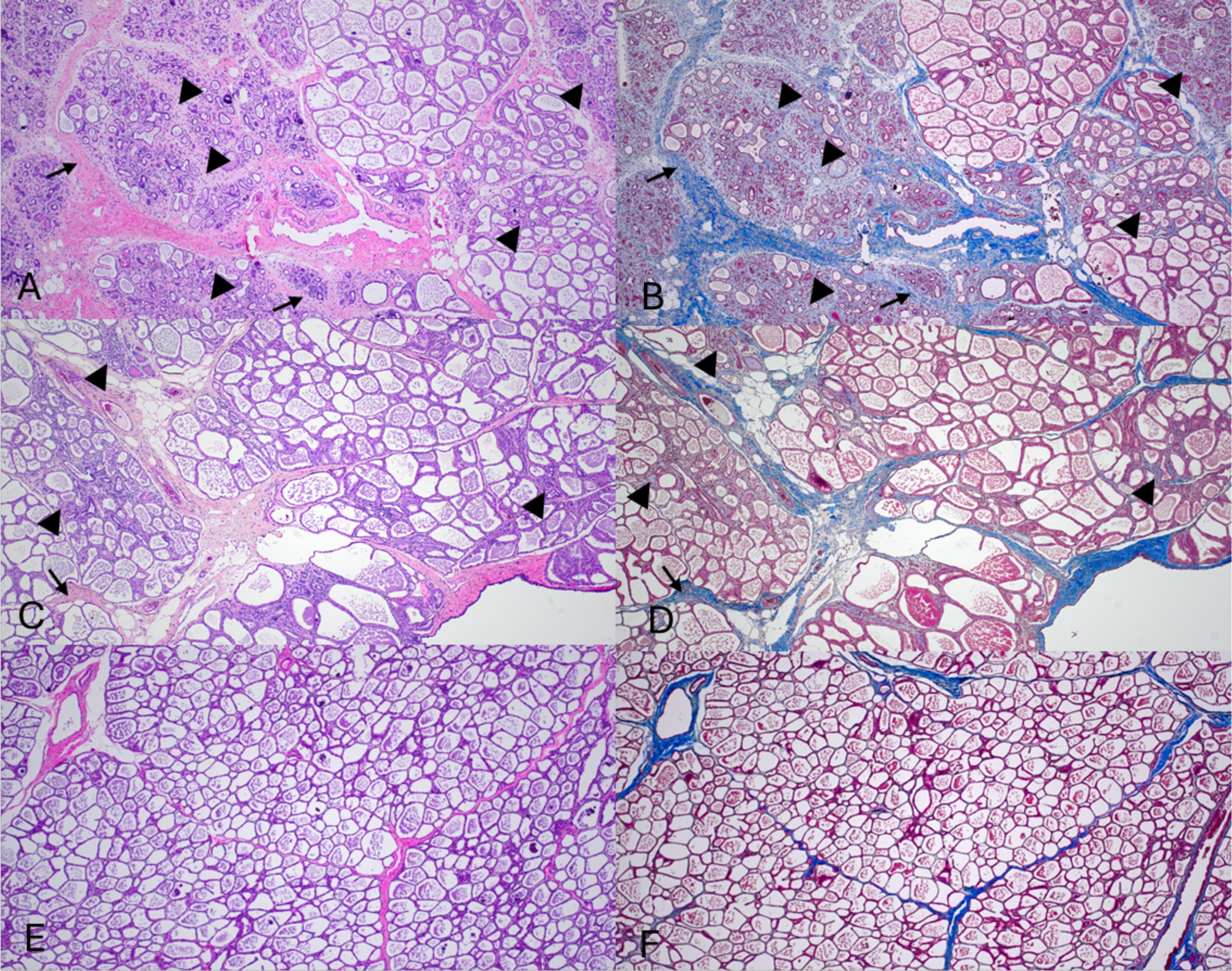
Interlobular (arrowheads) and intralobular (arrows) fibrosis of the left rear mammary gland in A) Cow 2112 and C) 2129. A matched Masson’s trichrome (fibrosis is blue) demonstrates the extent of fibrosis in B) Cow 2112 and D) 2129. A representative E) H&E and F) Masson’s trichrome in a non-inoculated teat (left front) of 2112 are provided for comparison. All photomicrographs are at 100X magnification.

IAV NP antigen was detected in the cytoplasm and nucleus of epithelial cells lining secretory alveoli in the inoculated mammary gland quarters of both cows at 24 DPI (Figure 6A-D). IAV NP antigen was also detected in the light zone of multiple germinal centers of the supramammary lymph node of cow 2129 (Supplemental Figure 2).

**Figure 6.**
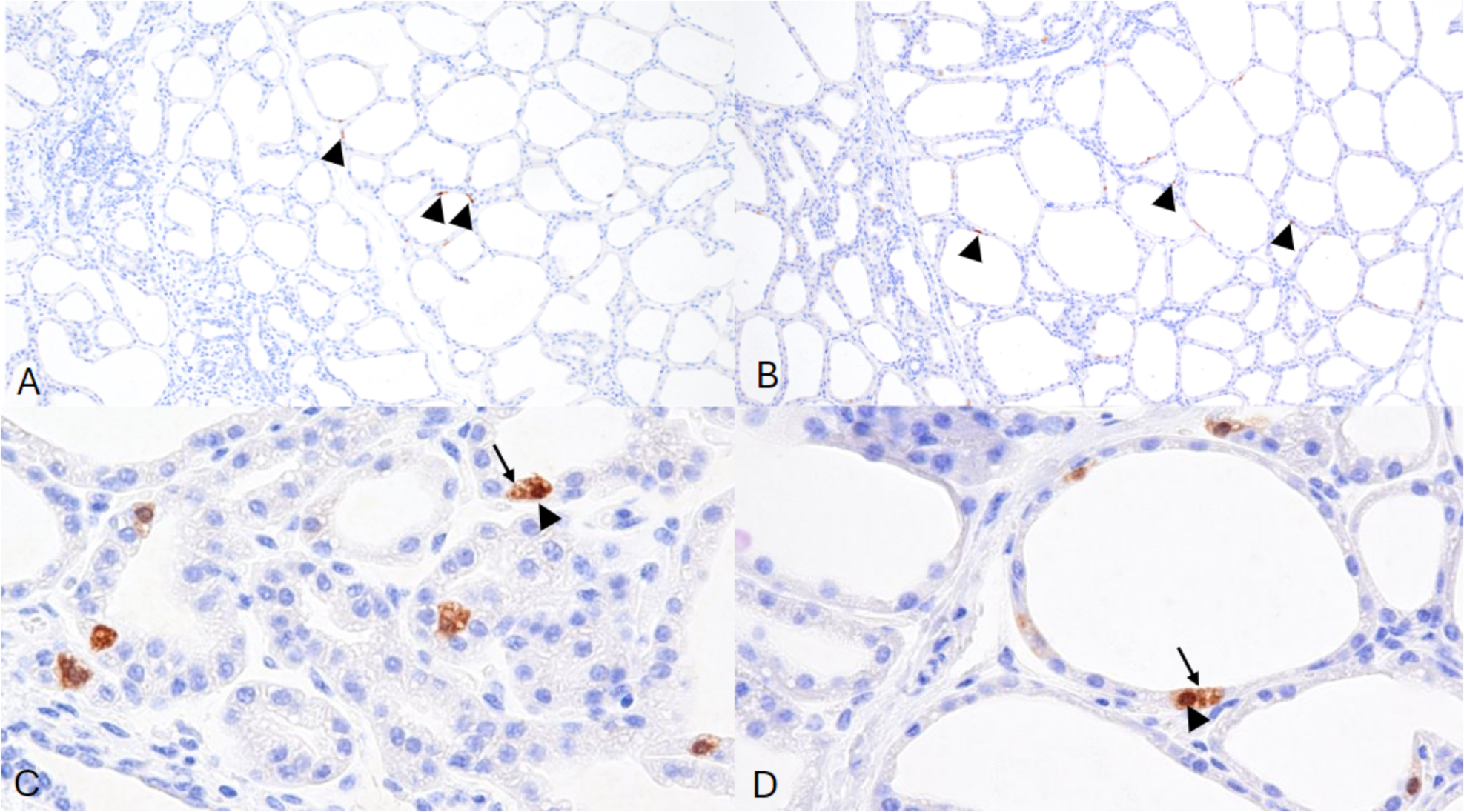
Influenza A virus antigen detection by immunohistochemistry (arrowheads) in the left rear mammary glands. Influenza A virus detection by immunohistochemistry in the cytoplasm (arrows) and nucleus (arrowheads) of epithelial cells lining secretory alveoli of the left rear mammary gland of A) Cow 2112 and B) Cow 2129. Photomicrographs at 100X (A and B) and 400X (C and D) magnification.

In the heifers necropsied at 7 DPI minimal lesions consistent with IAV infection were focused on conducting airways. The interstitium of a bronchiole of the caudal part of the left cranial lung lobe of Heifer 2311 was minimally circumferentially expanded by a predominance of lymphocytes and plasma cells. IAV was detected by IHC within the respiratory epithelial cells lining this conducting airway (Figure 7A-D). Bronchiolitis obliterans was present in the caudal part of the left cranial lung lobe of Heifer 2316 (Figure 7E). The lumen of a focal bronchiole was 80% occluded by a polyp of fibrocytes and fibrin admixed with inflammatory cells and lined by markedly attenuated epithelium. The surrounding interstitium was mildly circumferentially expanded by lymphocytes and plasma cells. Scant IAV NP antigen was detected in the remaining epithelial cells of the affected bronchiole, adjacent type II pneumocytes and probable alveolar macrophage (Figure 7F). Additional histologic lesions were observed in both cows and heifers; however, the role of IAV infection in the development of these lesions could not be defined by IHC or lesion character was not consistent with the disease time course.

**Figure 7.**
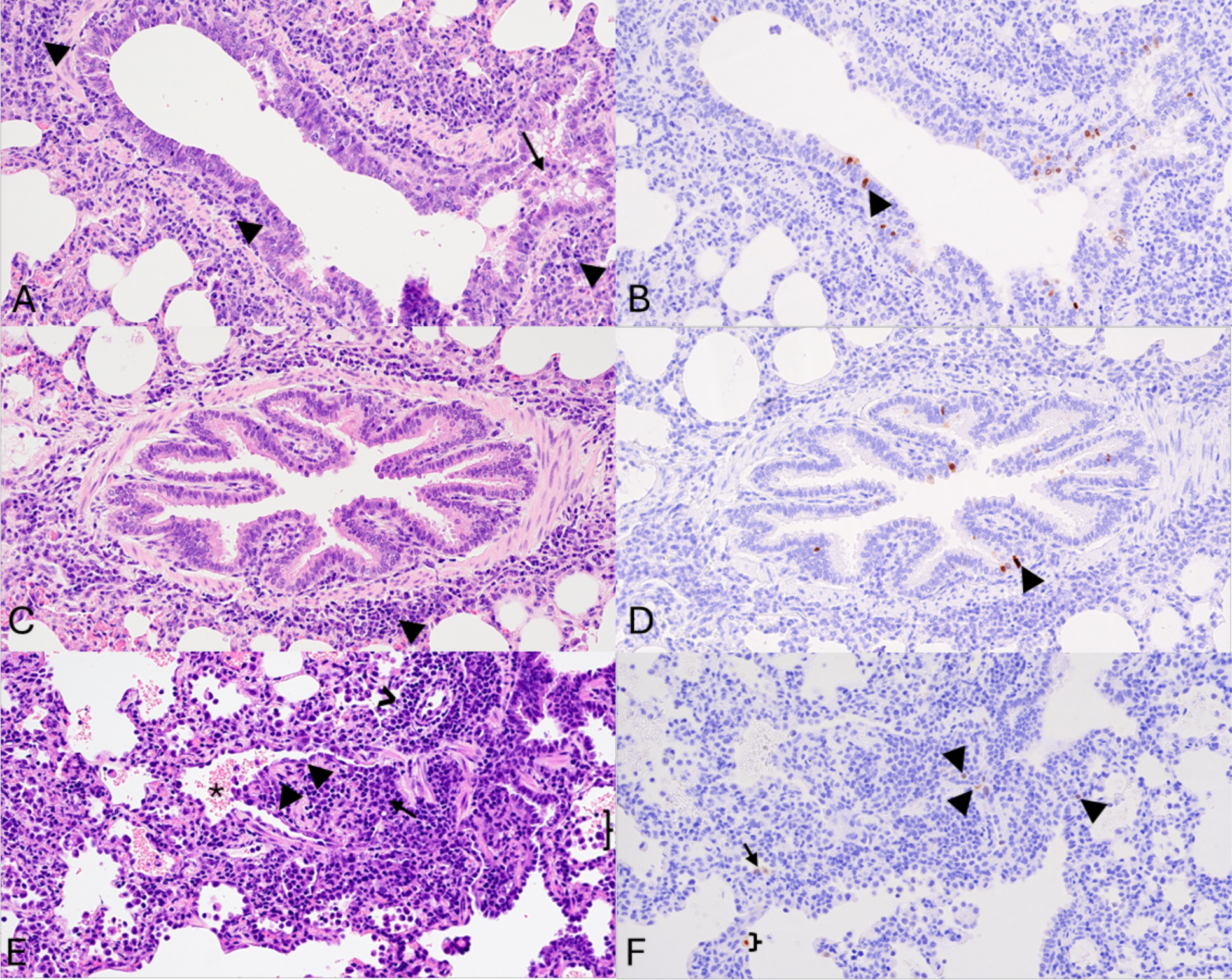
Peribronchiolar mononuclear cell inflammatory infiltrate (arrowheads) and accumulation of intraluminal inflammatory cells (arrow) in Heifer 2311 (A and C). Replication of highly pathogenic avian influenza virus in the respiratory epithelium lining the affected bronchiole of 2311 as demonstrated by immunohistochemistry (B and D; arrowheads). E) Bronchiolitis obliterans in Heifer 2316. The lumen of a bronchiole (asterisk) is partially occluded by a polyp lined by attenuated epithelial cells (arrowheads). The polyp is composed of fibroblasts, fibrin, and lymphocytes (arrow). Adjacent alveoli contain increased inflammatory cells (probable alveolar macrophages; brace). An adjacent arteriole is circumferentially surrounded by a mononuclear cell inflammatory infiltrate composed predominately of lymphocytes and fewer plasma cells (chevron). Replication of highly pathogenic avian influenza virus in the respiratory epithelium lining the affected bronchiole of 2316 as demonstrated by immunohistochemistry (F; arrowheads), probable type II pneumocytes (arrow) and alveolar macrophage (brace). All photomicrographs are at 200X magnification.

## Discussion

We describe the first two successful models of experimental infection in cattle with H5N1 clade 2.3.4.4b genotype B3.13 strain. Signs of clinical disease were variable and may not be recognized under field conditions, particularly from a respiratory route of exposure.

Experimentally inoculated cows showed clinical signs for 7-14 days, with changes to the milk similar to field reports, including change in color from white to yellow, thickening, and presence of flakes or clots ^2,30^. From 14-24 days after inoculation, after cows appeared to be recovering, viral RNA was still detected in the milk and replicating virus was present in the inoculated mammary gland quarters as detected by IHC on 24 DPI. However, viable virus was not found in the pooled milking machine bucket or individual quarter milk samples after 12 DPI. The detection of VN antibodies in the inoculated quarters coincided with negative virus isolation.

Additional functional antibody studies are needed for use with milk to understand when neutralizing antibodies appear and how long they last in larger numbers of animals. All calves and cows seroconverted during the study, confirming infection from both routes of inoculation. The commercial ELISA used to detect NP antibodies had not been validated by the manufacturer for bovine serum or milk at the time of this study. The kit cut-off value is 0.5 for avian species and included validation for H5N1 strains. The kit cut-off value is 0.6 for pigs but based on H1 and H3 subtypes. Robust validation with known positive and known negative bovine samples are needed for establishing a sensitive and specific cut-off for this host and subtype, but due to the known exposure, consecutive sampling, and steady decline of the S/N O.D. ratios, we used ≤0.6 as the cutoff for positivity.

The amount and duration of virus shed in milk from the inoculated mammary quarters are major findings of this study and point to the mammary gland and milk as primary sources of virus spread within and between dairy herds. This is consistent with the finding that movement of lactating cows was a primary epidemiologic link between the earliest herds involved in the outbreak^3^. Virus replication in infected mammary glands was considerably more than that expected from lungs of animals infected with host-adapted IAV. This is likely, at least in part, due to the cellular structure and physiology of the lung compared to the mammary gland. While IAV-susceptible epithelial cells line the conducting airways, these airways make up a minority of the lung structure. Based upon both NP IHC from naturally infected dairy cattle and lectin staining, most of the mammary gland is composed of IAV-susceptible epithelial cells ^2,31^. Moreover, there is regular sloughing of epithelial cells and secretion of milk fat globules. Upon release from the secretory epithelial cell, milk fat globules are bordered by epithelial cell membrane. Milk fat globules can also have a membrane crescent that consists of epithelial cell cytoplasm. Both the epithelial cell membrane and the crescent of milk fat globules may be lined by or contain non-infectious and/or infectious viral particles; however, further exploration of mammary tissue via electron microscopy is needed to confirm the role milk fat globules play in the shedding of HPAI.

The fibrosis and loss of secretory alveoli in the evaluated sections of the left rear mammary gland varied from mild to severe and were more extensive in sections evaluated in Cow 2112. The extent to which secretory alveoli were lost and replaced by fibrous connective tissue within the affected quarter or quarters may account for the variable return to milk production of individual animals following HPAI infection reported by farms ^2^. The long-term impact of mammary fibrosis in recovered cows in subsequent lactation cycles remains to be determined. Fibrosis is not a common finding in the lung following IAV infection and supports a divergent immunopathogenesis. This may be due to the need for limiting inflammatory responses in an organ that is vital to life, the number of cells susceptible to infection, and mechanisms of host viral clearance via immune responses. This hypothesis is further supported by the systemic clinical signs observed in the dairy cattle that included depression, anorexia, and drop in rumen motility. The presence of replicating virus in the inoculated mammary quarters at the time of necropsy twenty-four days after inoculation suggests a longer course for this organ to effectively clear the virus in comparison to the organs of the respiratory tract. Although fibrosis is a permanent change and would likely result in decreased milk production for animals in the field, prevention of fibrosis could be a primary variable for vaccine efficacy trials, along with significant reduction in virus titers. The application of a trichrome strain effectively demonstrated the amount of fibrous connective tissue present within glands and sections could be taken after the duration and quantity of viral shedding was characterized.

No to mild respiratory signs have been reported in affected dairy herds within the U.S. ^30^ and very few bovine respiratory diagnostic samples have been confirmed to be positive for H5N1 HPAI. The clinical signs, RT-qPCR results, macroscopic lesions, histologic findings, and antigen distribution of this study align with these field observations. However, the mild acute macroscopic and histologic lung lesions, detection of replicating IAV by IHC in the lower respiratory tract, and RT-qPCR with virus isolation in two upper respiratory swabs confirmed a respiratory phase of infection following aerosol inoculation. Although respiratory infection was limited in the 4 heifers, the detection of viable virus in 2 out of 4 represents a mode of infection and transmission that, when applied to an animal facility that commonly holds hundreds of animals, implies there is a role for the respiratory route. If natural exposure from the respiratory route leads to production of neutralizing antibodies, it may prove beneficial if subsequently re-exposed by the intramammary route during the milking process.

The enteric signs that included both diarrhea (Cow 2129) and dry fecal material (Cow 2112) noted in this study also align with reports from the field and justify the initial clinical differential of an atypical enteric bovine coronavirus infection for the milk drop syndrome. Minimal lesions were observed in the jejunum of both cows and cannot be definitively attributed to HPAI H5N1 infection. The loss of crypts and expansion of the lamina propria by fibrous connective tissue is a chronic change present at 24 DPI and due to study timing, tissues were collected days after enteric signs were noted. RT-qPCR did not detect HPAI in any rectal swab at any time point or the jejunum or feces of inoculated animals at necropsy, but low levels of replication within the upper gastrointestinal system cannot be ruled out based on timing of the necropsy. Additional studies focused on pathologic assessment and antigen distribution at multiple time points following inoculation should be conducted to more thoroughly evaluate tissue tropism at the cellular level.

The interspecies transmission of H5N1 clade 2.3.4.4b to many mammal species, now including cattle with genotype B3.13, is unprecedented in our understanding of avian-adapted IAV ^32,33^. This raises concern for other mammalian hosts, including pigs and other domestic livestock and pets, and particularly for humans. The human cases in the United States have been clinically mild and limited in number ^4,5^, but concern remains as the H5N1 continues to expand into new hosts, spread geographically, and reassorts with other avian or mammalian subtypes. The possibility of H5N1 becoming endemic in cattle increases as the number of infected herds continues to rise ^1^. Pasteurization was shown to inactivate virus and retail milk remains negative for infectious virus, thus not a risk for human consumption when processed according to Food and Drug Administration standards ^34^. Unpasteurized milk and dairy products are a risk to humans and other animals. Milk diverted from the human food supply in H5N1 positive dairy herds or from suspect cows should not be fed to other farm or peridomestic animals. The sustained transmission among dairy cattle is an animal health crisis due to production and economic losses and is a public health challenge due to occupational exposure on dairy farms. The development of reproducible experimental challenge models like the ones described here is the essential first step to inform subsequent research on intervention and vaccination strategies. Although limited in the number of animals due to their size and high containment space requirements, we reproduced the clinical observations from the field of viral mastitis due to HPAI H5N1 infection alone and confirmed respiratory involvement. Further studies to understand transmission, refine the pathogenesis model, and define the kinetics of protective immunity in cattle infected with HPAI are urgently needed.

## Data Availability

Data that support the findings of the clinical challenge studies and sequence data, code, and materials used in the analysis are available at https://github.com/flu-crew/datasets. Sequence data generated within this study are provided at NCBI GenBank and the accession numbers are provided in the supplementary materials.

## Acknowledgments

We thank the USDA NADC and NVSL leadership and personnel from animal resource and facilities and engineering units and the NVSL Diagnostic Virology Laboratory, without whom the study could not have been successfully conducted. Katharine Young, Emily Love, Sarah Anderson, Dan Coster, Laura Kepler, Sebastian O’Bryon, Seth Van Dexter, Josh Thompson, Josiah Fichter, and Sydney Smith are recognized for laboratory technical assistance and Tonia McNunn and the NCAH select agent staff for compliance assistance. We thank Dr. Rebecca Cox, Jason Huegel, Tiffany Williams, Jonathan Gardner, Jared Peterson, Emma Hay, Emma Pratt, and Brian Conrad for assistance with animal studies. We thank Drs. Paul Plummer, Phillip Jardon, and Luis Gimenez-Lirola from the Iowa State University Veterinary Diagnostic and Production Animal Medicine department for helpful discussions on the study and milking equipment. This work was supported in part by the U.S. Department of Agriculture (USDA) Agricultural Research Service (ARS project number 5030-32000-231-000-D); the National Institute of Allergy and Infectious Diseases, National Institutes of Health, Department of Health and Human Services (contract numbers 75N93021C00015); and the SCINet project and the AI Center of Excellence of the USDA Agricultural Research Service (ARS project numbers 0201-88888-003-000D and 0201-88888-002-000D). Mention of trade names or commercial products in this article is solely for the purpose of providing specific information and does not imply recommendation or endorsement by the U.S. Government. USDA is an equal opportunity provider and employer.

## Supplemental Files

**Supplemental Table 1.** Clinical scores used to assess dairy calves and lactating dairy cows*.

| *Score* | *Behavior* | *Respiratory signs/Rate* | *Cough* | *Nasal discharge* | *Ocular Discharge* | *Feces* |
| --- | --- | --- | --- | --- | --- | --- |
| 0 | Normal | Normal | Normal | Normal, serous discharge | Normal | Normal |
| 1 | Mild lethargy with decrease in ambulation and attitude compared to pen mates | Slightly increased respiratory effort/slight dyspnea | Induce single cough | Small amount of unilateral, cloudy discharge | Mild ocular discharge (predominately in canthi) | Semi-formed, pasty |
| 2 | Moderate lethargy with stimulation needed to provoke ambulation | Notable increase in respiratory effort/dyspnea | Induce repeated coughs or occasional spontaneous cough | Cloudy or excessive mucus | Moderate bilateral ocular discharge | Loose, but stays on top of bedding or dry and tacky |
| 3 | Marked lethargy with ambulation not provoked by stimulation | Severe dyspnea with respiratory distress and tachypnea | Repeated spontaneous coughing | Copious, mucopurulent nasal discharge | Heavy ocular discharge (lashes coated) swelling/erythema of the eyelid | Watery, sifts through bedding |

\*Adapted from https://www.vetmed.wisc.edu/fapm/svm-dairy-apps/calf-health-scorer-chs/ Calf Health Scorer (CHS) – Food Animal Production Medicine – UW–Madison ([wisc.edu](https://www.vetmed.wisc.edu/fapm/svm-dairy-apps/calf-health-scorer-chs/))

**Supplemental Table 2.**
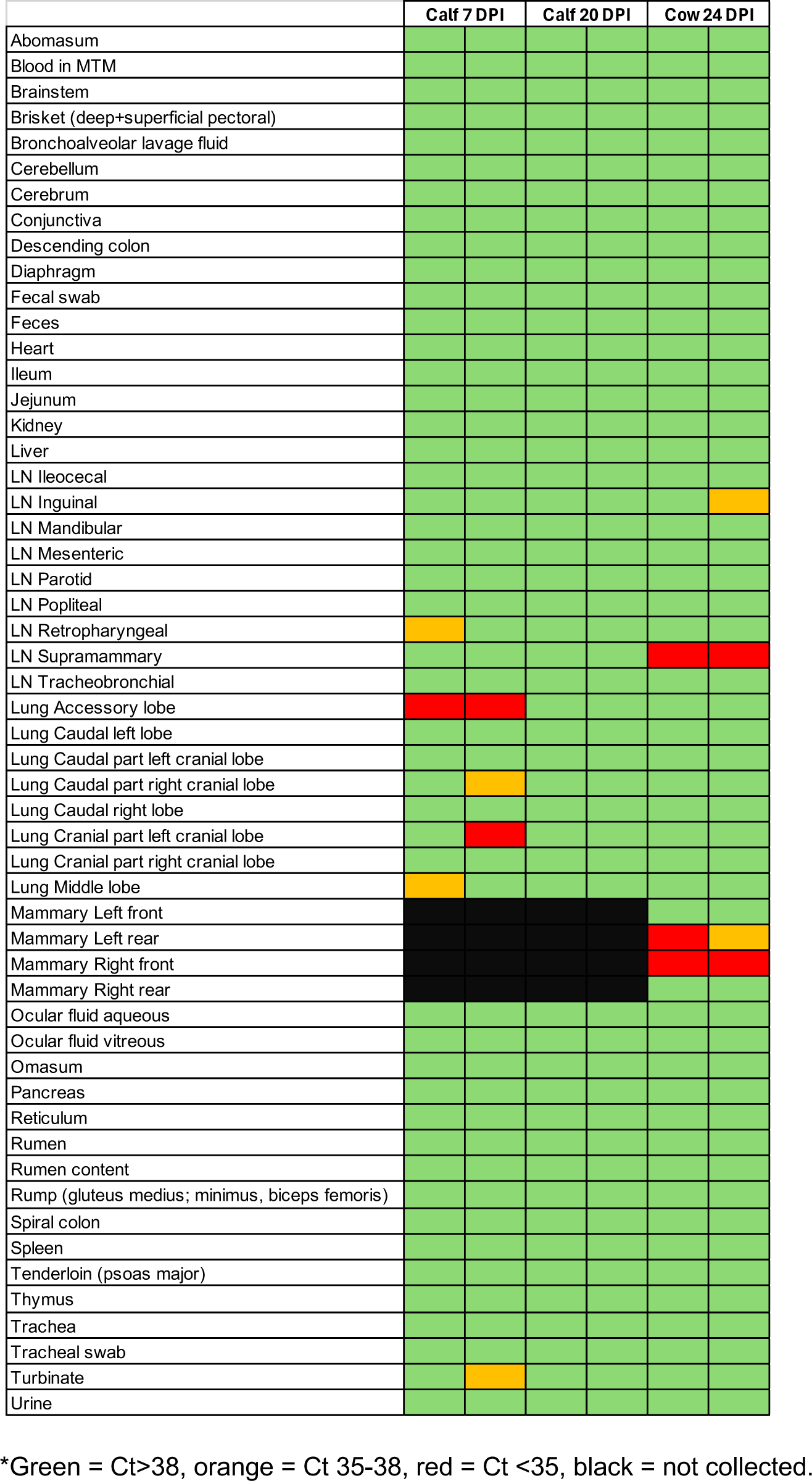
Tissues and samples collected at necropsy and tested by RT-qPCR.*

**Supplemental Table 3.** Positive RT-qPCR samples in aerosol inoculated heifer calves.*

|  | DPI | 1 | 2 | 3 | 4 | 5 | 6 | 7 | CUMULATIVE |
| --- | --- | --- | --- | --- | --- | --- | --- | --- | --- |
| Nasal swab |  | 1<br>(32.3)<sup>#</sup> | 1<br>(24.8) | 1<br>(30.2) | 1<br>(31.6) | 1<br>(27.5) | 1<br>(36.2) | 1<br>(29.8) | 7 |
| Oropharyngeal swab (FLOQ) |  | 1<br>(37.3) | 1<br>(34.9) | 2<br>(34) | 1<br>(36.8) | 0 | 0 | 0 | 5 |
| Oropharyngeal swab (COTT)* |  | 1<br>(37.9) | 3<br>(31.6) | 0 | 1<br>(36.3) | 0 | 1<br>(33.4) | 0 | 6 |
| Ocular swab |  | 0 | 3<br>(36.8) | 0 | 0 | 1<br>(31.4) | 0 | 1<br>(35.5) | 5 |
| Saliva |  | 0 | 1<br>(34.2) | 2<br>(35.3) | 1<br>(33.5) | 1<br>(35.3) | 1<br>(37.4) | 0 | 6 |
|  | CUMULATIVE | 3 | 9 | 5 | 4 | 3 | 3 | 2 |  |

<sup>#</sup>Number of samples with Ct<38 out of four and RT-qPCR Ct value or average in parentheses.
\*Virus isolation was unsuccessful for attempted cotton oropharyngeal swabs. Five FLOQ nasal swabs from 2316 and one FLOQ oropharyngeal swab from 2311 were positive by virus isolation.

**Supplemental Table 4.**
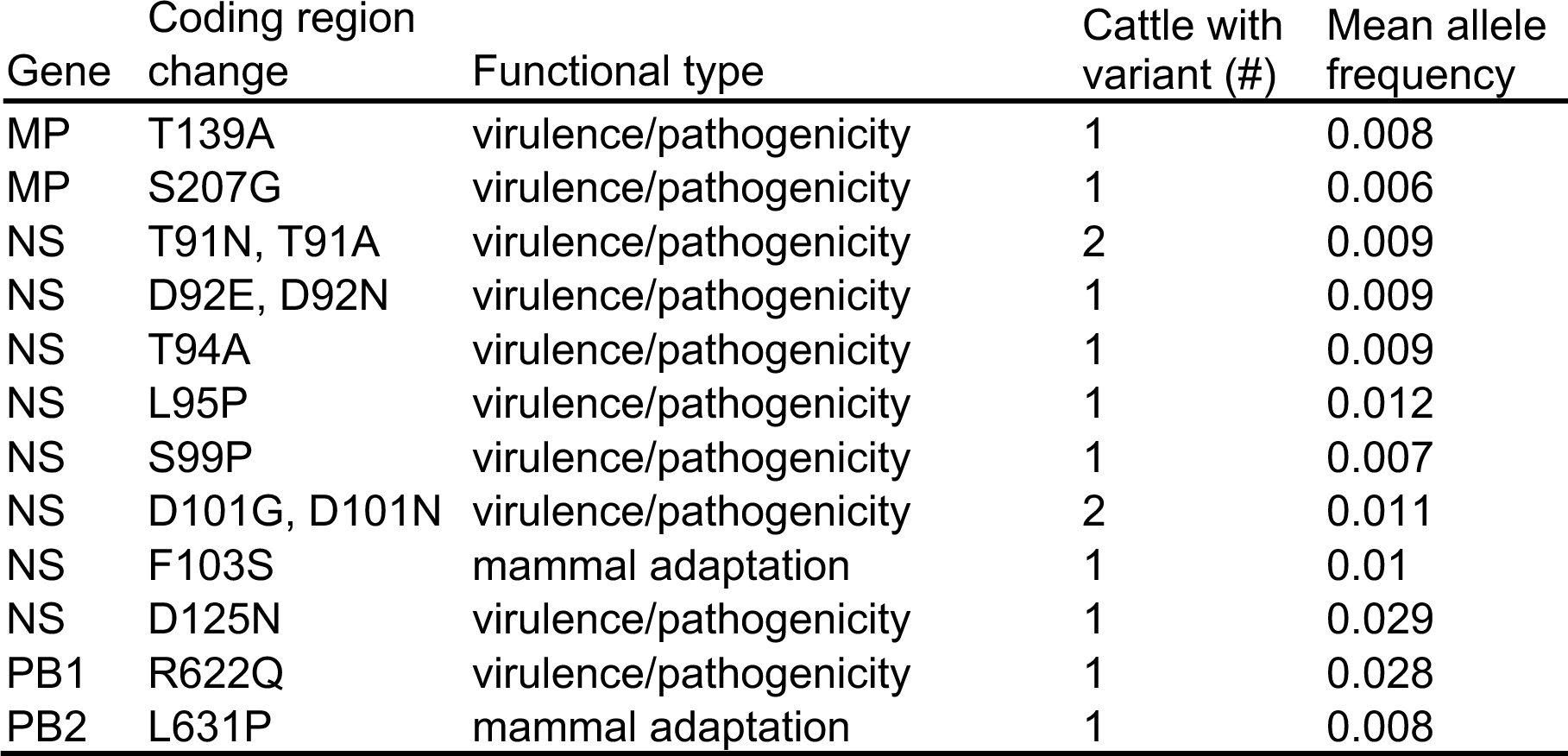
Raw read data collected from cattle were processed and high- and low-frequency single nucleotide variants (SNVs) were identified. The SNVs that resulted in a coding region change were screened against known functional changes with a relevant selection included here. No SNVs from this database were detected in the consensus sequence and remained at low frequencies. The number of cattle with the SNV were counted and mean allele frequency was calculated.

**Supplemental Figure 1.**
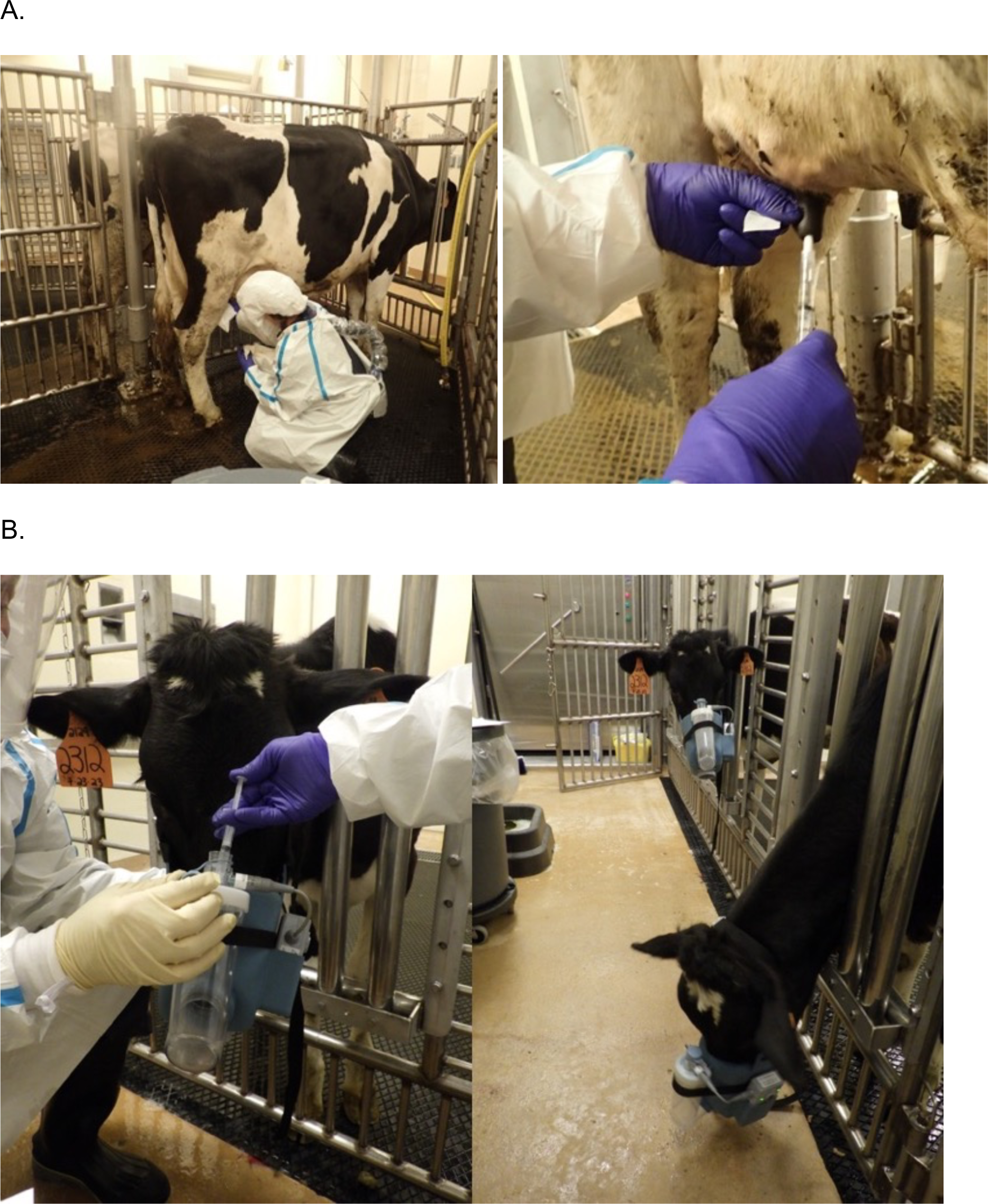
A) Photographs of intramammary inoculation in the lactating Holstein cows via teat canula. B) Photographs of aerosol respiratory inoculation in the Holstein heifer calves using nebulizer masks.

**Supplemental Figure 2.**
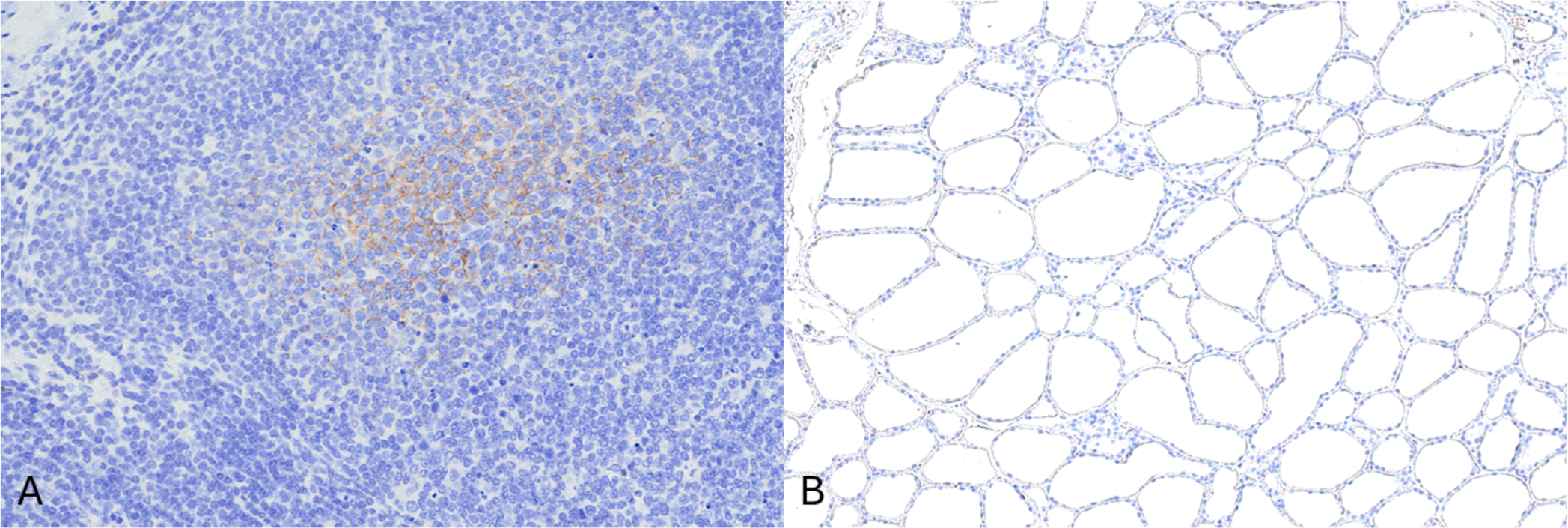
Influenza A virus detection by immunohistochemistry in the germinal center light zone of the supramammary lymph node of cow 2129 (A). Immunohistochemistry did not detect Influenza A virus antigen in the left front mammary gland of 2112 (C). Photomicrographs at 200X (A) and 100X (B) magnification

